# Population genomic analysis reveals contrasting demographic changes of two closely related dolphin species in the last glacial

**DOI:** 10.1101/225987

**Authors:** Nagarjun Vijay, Chungoo Park, Jooseong Oh, Soyeong Jin, Elizabeth Kern, Hyun Woo Kim, Jianzhi Zhang, Joong-Ki Park

## Abstract

Population genomic data can be used to infer historical effective population sizes (*N*_e_), which help study the impact of past climate changes on biodiversity. Previous genome sequencing of one individual of the common bottlenose dolphin *Tursiops truncatus* revealed an unusual, sharp rise in *N*_e_ during the last glacial, raising questions about the reliability, generality, underlying cause, and biological implication of this finding. Here we first verify this result by additional sampling of *T. truncatus.* We then sequence and analyze the genomes of its close relative, the Indo-Pacific bottlenose dolphin *T. aduncus.* The two species exhibit contrasting demographic changes in the last glacial, likely through actual changes in population size and/or alterations in the level of gene flow among populations. Our findings demonstrate that even closely related species can have drastically different responses to climatic changes, making predicting the fate of individual species in the ongoing global warming a serious challenge.

---

Predicting the impact of a rapidly changing climate on biodiversity represents an urgent challenge due to the ongoing global warming (Thuiller 2007). Although the causes of past and present climatic changes may be different, the consequences of past climatic changes on biodiversity can provide useful insights. Hence, a promising approach is to use population genomic data to infer historical effective population sizes (*N*_e_) and study how *N*_e_ has responded to past climatic change events such as Pleistocene glacial cycles. This approach has frequently revealed a reduction in *N*_e_ during the last glacial (110-12 kya) for temperate species, including marine cetaceans (Moura et al. 2014; Yim et al. 2014). Surprisingly, however, the genomic analysis of one individual of the common bottlenose dolphin *Tursiops truncatus* suggested a steep rise in *N*_e_ in the last glacial (Yim et al. 2014), raising questions about the reliability, generality, underlying cause, and biological implication of this finding. In the present study, we first verify the previous finding by additional sampling of *T. truncatus*. We then sequence and analyze the genomes of its closest congener, the Indo-Pacific bottlenose dolphin *T. aduncus*, for comparison. We report contrasting demographic changes of these two species in the last glacial, explore potential causes, and discuss implications.

Bottlenose dolphins (genus *Tursiops)* include at least two taxonomically accepted, widespread species *(T. truncatus* and *T. aduncus)* that diverged from each other ~2.5 Mya (Vilstrup et al. 2011). They occupy distinct but overlapping habitats and regions (Hale et al. 2000): *T. truncatus* is distributed in the coastal, near-shore, and off-shore zones of most oceans worldwide, with the exception of polar areas, whereas *T. aduncus* is discontinuously distributed along the coastal areas of warm temperate and tropical Indian and Indo-Pacific oceans, from east Africa to the northwestern Pacific (**Fig. 1**).

**Figure 1.**
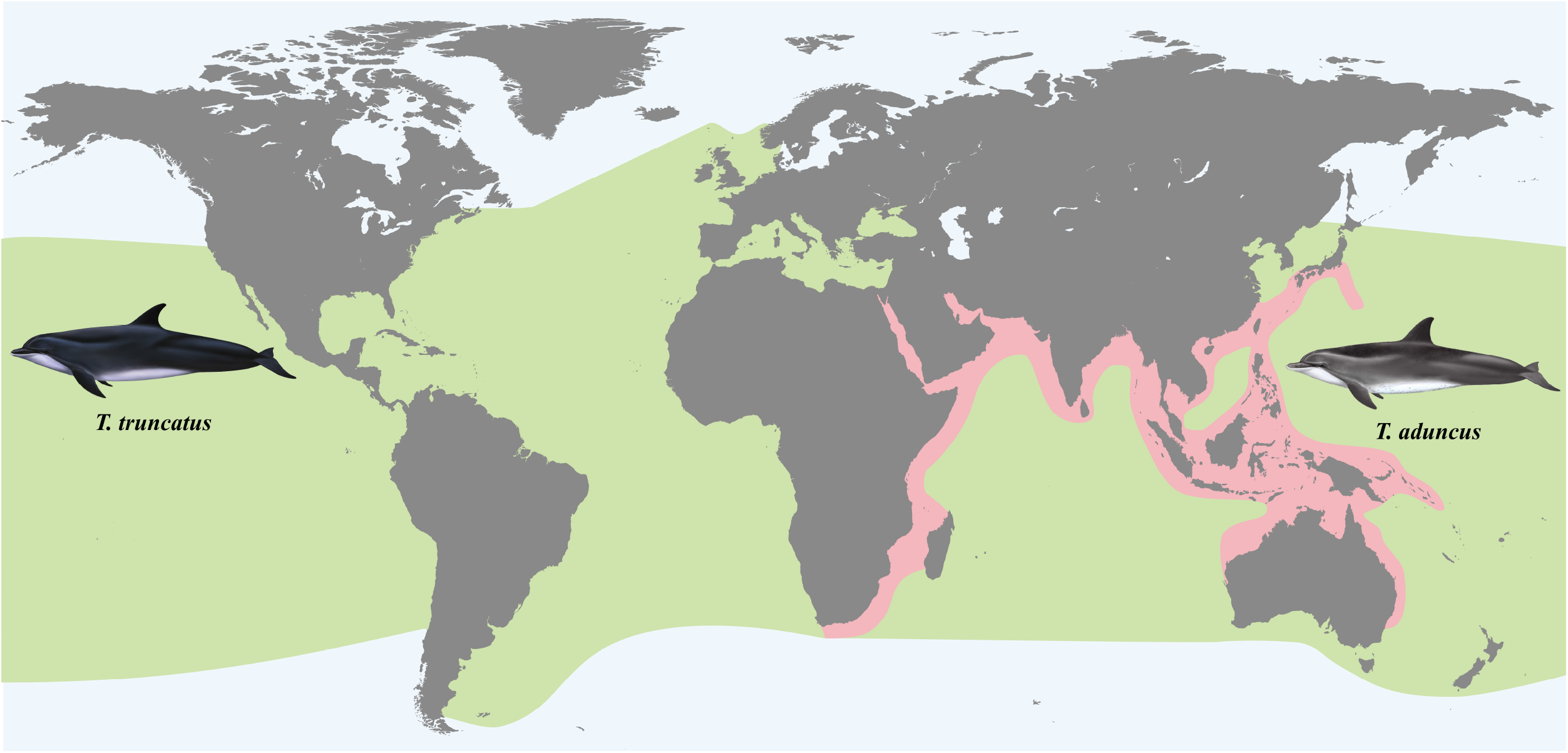
Map showing the geographic distributions of the common bottlenose dolphin *Tursiops truncatus* (green) and Indo-Pacific bottlenose dolphin *T. aduncus* (pink), based on information from the Whale and Dolphin Conservation Society.

We used the Pairwise Sequentially Markovian Coalescent (PSMC) method (see Materials and Methods) to respectively infer the temporal changes in *N*_e_ from publically available genome sequences of four *T. truncatus* individuals, including the one previously analyzed (Yim et al. 2014) (**Table S1**). For all four genomes, *N*_e_ rose sharply starting around the beginning of the last glacial, peaked when the atmosphere temperature reached the minimum, and then dropped as the temperature rebounded (**Fig. 2**). Bootstrap analysis confirms the statistical robustness of these trends (**Fig. S1A**).

**Figure 2.**
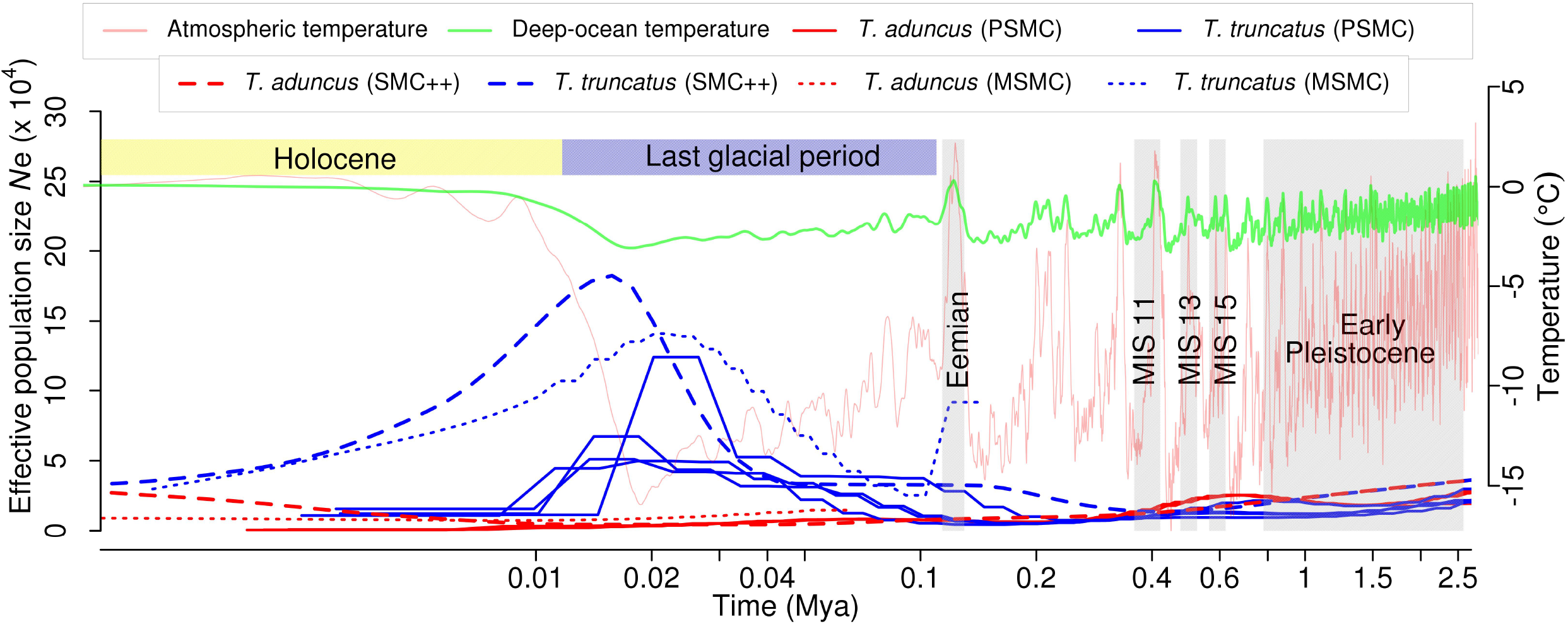
Contrasting demographic changes of the two bottlenose dolphin species in the last glacial, inferred from genome sequences using PSMC, MSMC, and SMC++. MIS, Marine Isotope Stage.

To probe whether this unusual demographic pattern is specific to *T. truncatus*, we generated a high-quality *de novo* genome assembly for *T. aduncus,* using 180× coverage of Illumina sequencing of an individual sampled from Korea (**Table S2**), followed by sequencing of three additional individuals at lower coverages (22-32×) (see Materials and Methods). Population genomic analyses were conducted using four *T. truncatus* and four *T. aduncus* individuals. Nucleotide diversity (π) is greater for *T. truncatus* (0.0015) than *T. aduncus* (0.00095), as expected from the wider geographic distribution of the former than the latter. The two species exhibit high levels of genome-wide differentiation (mean *F*_st_ = 0.61), consistent with their different geographic distributions and classification as distinct species (with the caveat that this conclusion is based on East Asian *T. truncatus*; see below). The PSMC analysis of each *T. aduncus* individual shows a decline in *N*_e_ during the last glacial (**Fig. 2, Fig. S1A**). This prolonged decline started more than 0.5 Mya, and no rebound since then is apparent (**Fig. 2**). The PSMC-inferred, contrasting patterns of *N*_e_ between the two dolphin species are robust to different generation times and mutation rates assumed (**Fig. S1B**) and are confirmed by MSMC and SMC++ (**Fig. 2**), which are improved methods that can simultaneously analyze multiple genome sequences (see Materials and Methods). The population size trajectories that have not been scaled by mutation rate and generation time also show a clear distinction between the two species (**Fig. S1C**).

All individuals of *T. truncatus* and *T. aduncus* analyzed above were sampled from East Asia (China, Korea, and Japan) and are thus more or less comparable. The public domain also houses the genome sequences from two individuals of *T. truncatus* sampled off Isle au Pitre in the Mississippi Sound, Louisiana and the coastal waters of the US eastern seaboard, respectively. As expected, multidimensional scaling (**Fig. S2A**) and principal component analysis (**Fig. S2B**) both show that all *T. truncatus* individuals are genetically distinct from *T. aduncus.* The four *T. aduncus* individuals are genetically highly similar (**Fig. S2**). Within *T. truncatus*, however, the two American individuals are genetically quite distinct from each other and from the four East Asian individuals (**Fig. S2**), suggesting population differentiation of *T. truncatus.* Strikingly, the Louisiana individual has a temporal trend of *N*_e_ highly similar to those of *T. aduncus* individuals, with a decline in *N*_e_ in the last glacial (**Fig. S3**). The genome sequence coverage of the US eastern seaboard individual (0.7×) is too low to allow a reliable PSMC analysis.

Because the tremendous temporal changes of *N*_e_ for (East Asian) *T. truncatus* coincided well with the drastic atmosphere and deep-ocean temperature changes of the last glacial (**Fig. 2**), the demographic changes were likely caused directly or indirectly by the climatic changes. Global cooling could directly affect dolphin habitats and distributions because of lowered sea levels and reduced ocean temperatures (Learmonth et al., 2006). While these effects could explain the *N*_e_ decline for the coastal species *T. aduncus*, they cannot easily explain the *N*_e_ increase for the broadly distributed *T. truncatus*. Global climate change could have also affected dolphins through food availability due to alterations in currents, upwelling, and productivity, and through the altered ecology of pathogens, competitors, and predators (Hoegh-Guldberg and Bruno 2010). It is notable that the *N*_e_’s of some dolphin predators declined during the last glacial, which could lead to a surge in dolphin *N*_e_. For instance, dolphins are preyed upon by some large sharks, most of which are restricted to the warmer tropical climates. In general, sharks in warm zones show genetic signatures of reduced *N*_e_ in the last glacial (O’Brien et al. 2013). Furthermore, killer whales show a decline in *N*_e_ at the same period (Moura et al. 2014). Because killer whales are more abundant in *T. truncatus*’ habitat than *T. aduncus*’ habitat (Forney and Wade 2007), *T. truncatus* may have benefited more than *T. aduncus* from killer whale’s decline in the last glacial. While dolphins are a relatively minor part of the diet of killer whales today, it is possible that climate changes indirectly caused the contrasting trends in *N*_e_ between *T. truncatus* and *T. aduncus* by differentially impacting their predators.

Temporal changes in *N*_e_ can also be caused by alterations in population structure or gene-flow between populations (Mazet et al. 2016). Marine mammals with their complex patterns of ancestry can especially be influenced by such processes (Foote and Morin 2016). To examine this possibility in the dolphins, we conducted a PSMC analysis of pseudo-diploids artificially constructed from genomes of different individuals (see Materials and Methods). Pseudo-diploids made from different *T. aduncus* individuals and actual *T. aduncus* individuals show almost identical temporal trends of *N*_e_ (**Fig. 3**), confirming a lack of population structure in this species. For (East Asian) *T. truncatus*, however, all pseudo-diploids show an increase in *N*_e_ towards infinity from 110 to 40 kya during the last glacial (**Fig. 3**), indicating the presence of population structure, echoing a previous genetic marker-based study that reported population structure of *T. truncatus* in the North-East Atlantic despite no obvious barrier to gene flow (Louis et al. 2014). SMC++ split analysis using the joint frequency spectrum confirms the PSMC results of pseudodiploids by showing much earlier split times between (East Asian) *T. truncatus* individuals than between *T. aduncus* individuals (**Fig. S4**). Our results, coupled with the hump in *N*_e_ from the real *T. truncatus* individuals during the last glacial (**Fig. 2**), suggest a reduction in gene flow among *T. truncatus* populations in this period (Mazet, et al. 2016), which was probably caused directly or indirectly by the global cooling. In line with the population structure inferred from the multidimensional scaling (**Fig. S2A**) and principal component analysis (**Fig. S2B**), PSMC analysis shows earlier rises in *N*_e_ to infinity for pseudo-diploids constructed between Louisiana and East Asian *T. truncatus* individuals than for pseudo-diploids between East Asian individuals (**Fig. S5**).

**Figure 3.**
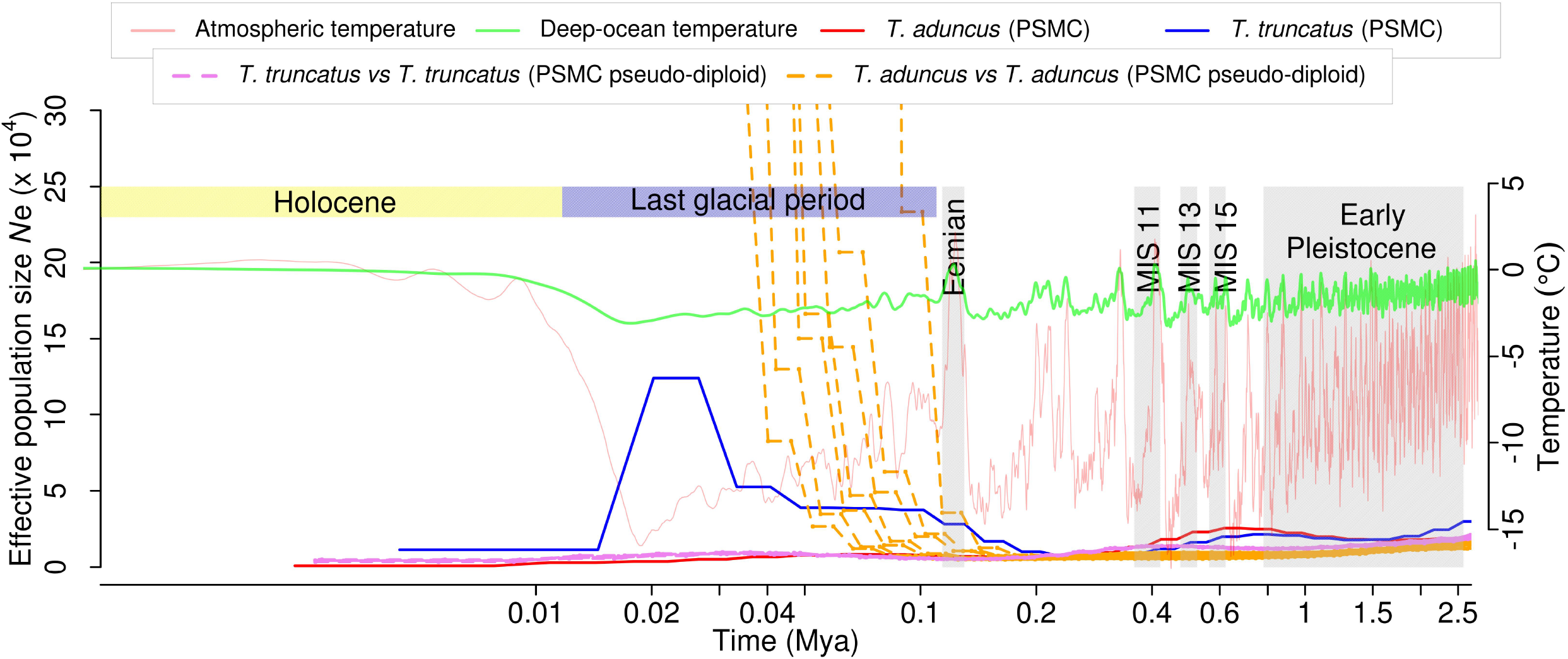
Pseudo-diploid analysis using PSMC suggests population structures in (East Asian) *T. truncatus* but not *T. aduncus.*

In summary, our population genomic analysis revealed contrasting temporal trends of *N*_e_ in the last glacial between two closely related dolphin species and potentially even between populations within a single species, likely due to complex and often idiosyncratic ecological interactions that vary between species or populations, including for example changes in predator population sizes and migration rates among populations. Such variations make it possible for closely related species and populations to respond drastically differently to the same climate event. The pattern reported here is unlikely to be unique to bottlenose dolphins, because similar contrasts were previously inferred for sea snails (Albaina et al. 2012) and beltfish (He et al. 2014) on the basis of much smaller data and in minke whales on the basis of genomic data of fewer samples (Kishida 2017). Hence, predicting the impact of climate change on a particular species or population is likely difficult without a much greater understanding of the specific ecological and biological factors involved.

## MATERIALS AND METHODS

### Dolphin samples

*T. aduncus* and *T. truncatus* were considered monospecific until the recognition of two species on the basis of genetic evidence and morphology (osteology and external morphology including beak morphology, the shape of dorsal fin, and the presence/absence of ventral spotting in adults) (Wang et al. 1999; Wang et al. 2000; Wang et al. 2000). More recently, molecular evidence suggests that *T. aduncus,* long-beaked common dolphin *Delphinus capensis,* and striped dolphin *Stenella coeruleoalba* form a monophyletic clade that is sister to *T. truncatus* (Leduc et al. 1999; Vilstrup et al. 2011; Moura et al. 2013).

Tissue samples of four individuals of *T. aduncus* were obtained from stranded dead individuals on the coast of Jeju Island, Korea. Sex was recorded in the field, and confirmed by subsequent genetic analysis. We generated whole-genome sequencing data at ~180× coverage (based on an estimated 2.29 Gb genome size; **Table S1**) for one individual and re-sequencing data of 22–32× coverage for the remaining three individuals (**Table S1**). These data were supplemented with publicly available sequences of six individuals of *T. truncatus* (four from East Asia and two from the US) downloaded from the Short Read Archive (SRA). Details of the coverage, sex, and SRA accession numbers of all samples are provided in **Table S1**.

### Genomic DNA library construction, sequencing, and assembly

Genomic DNA was extracted from muscle tissues using the DNeasy Blood & Tissue kit (Qiagen, Germany). Extracted DNA was quantified by the Quant-iT BR assay kit (Invitrogen). For the reference genome of *T. aduncus,* we constructed short-insert (350 and 550 bp with 2×251 bp reads) and long-insert (5 and 10 kb with 2×101 bp reads) libraries using the standard protocol provided by Illumina (San Diego, USA). Three additional *T. aduncus* individuals were subjected to 2×150 bp paired-end sequencing of a 350 bp insert library. The raw reads were pre-processed using Trimmomatic v0.33 (Bolger et al., 2014) and Trim Galore (Martin, 2011), in which reads containing adapter sequences, poly-N sequences, or low-quality bases (below a mean Phred score of 20) were removed. For the reference genome assembly of *T. aduncus*, we used all preprocessed reads from four (350 bp, 550 bp, 5 kb, and 10 kb) libraries and employed ALLPATHS-LG v52488 (Gnerre et al. 2011). Next, the gaps (any nucleotide represented by “N” in scaffolds) were closed using GapCloser, a module of SOAPdenovo2 (Luo et al. 2012). For further analyses, any scaffold with > 0.04% hits belonging to bacterial genomes from NCBI (release of Nov. 2016), downloaded from the RefSeq microbial genomes ftp site (Tatusova et al. 2014), was removed. A draft genome was assembled for *T. aduncus* using both paired-end and mate-pair libraries. The final assembled genome is 2.5 Gb in total length, close to the estimate by K-mer analysis (2.29 Gb), and was composed of 16,188 scaffolds (≥ 1 kb in their length) with an N50 value of 1,254 kb (Table S2). The assembly was found to be of good quality; 95.2% of the core eukaryotic genes (based on 248 core essential genes) from CEGMA v2.5 (Parra et al. 2007), 89.7% of the core vertebrate genes (based on 233 one-to-one orthologs in vertebrate genomes) previously identified (Hara et al. 2015), and 92.1% of the metazoan single-copy orthologs (based on 843 genes) from BUSCO v1.22 (Simão et al. 2015) could be identified in the genome assembly.

### Read mapping and variant calling

The genome assembly and annotation of *T. truncatus* release-87 were downloaded from the Ensembl. The sequencing reads from each dolphin species were mapped to the genome assembly using the Burrows-Wheeler aligner BWA-MEM (Li 2013), with default settings. PCR duplicate reads were collapsed using the rmdup option in SAMtools (v 0.1.19) (Li et al. 2009). Regions of the genome annotated as repeats by RepeatMasker (Smit et al. 1996) or mitochondrial DNA transposed to the nuclear genome (numts) identified by blast hits to the mitochondrial sequence (at an E-value threshold of 10^−3^) were removed from further analysis. We identified variable sites across the genome using the ‘mpileup’ command in samtools.

### Identification of sex chromosome markers and sexing of individuals

We searched for the previously identified Y chromosome marker from the SRY gene (Palsbøll et al. 1992) in sequencing reads to identify/validate the sex of each individual. The inferred sex always matched the previously recorded sex in the field. The scaffolds of the genome assembly from a female *T. truncatus* individual were grouped into autosomal and X chromosome scaffolds based on the ratio of sequencing coverage between female and male individuals, using a previously described approach (Bidon et al. 2015). Briefly, the mean coverage of all scaffolds longer than 200 kb was calculated using BEDTools (Quinlan and Hall 2010). Scaffolds that consistently showed a female/male coverage ratio near two were deemed to be from the X chromosome. As a negative control, we also examined the coverage ratio of male/male and female/female individuals.

### Demographic history and population structure

Reads mapped to autosomal scaffolds longer than 100 kb were analyzed using the program PSMC (Li and Durbin 2011) to investigate temporal trends of Ne, under the assumption of a generation time of 21 years and a mutation rate of 1.21×10^−9^ per site per year for both dolphin species (Taylor et al. 2007; Yim et al. 2014). It is recognized that using ~75% of the genome or more in the PSMC analysis results in reliable results (Nadachowska-Brzyska et al. 2016). Our PSMC analysis used 74% of the assembled genome after all the filtering had been performed. Another common source of error in the PSMC analysis is the use of low-coverage genome sequences. By down-sampling a high-coverage genome sequence to various levels of coverage, Nadachowska-Brzyska et al. showed that results start to differ from the original result when the coverage is below 20× (Nadachowska-Brzyska et al. 2016). However, whether this finding is general or specific to the particular species/genome used is unclear. In our study, we generated whole genome sequencing data from four individuals of *T. aduncus,* with 22-180× coverages (**Table S1**), and all four individuals show similar *N*_e_ plots from the PSMC analysis. We used publicly available datasets from six *T. truncatus* individuals in this study, including four from East Asia and two from the US. Among the four East Asian samples, one has a 43× coverage and was previously analyzed using PSMC (Yim et al. 2014). Our reanalysis of this sample using PSMC yielded similar results. Three additional East Asian samples have 10-12× coverages and may not be ideal for the PSMC analysis. Nevertheless, we found the *N*_e_ plots from these three individuals highly similar to that of the 43× coverage individual. Hence, the relatively low coverage does not seem to affect the PSMC analysis in the present case. Of the two US samples, one has < 1× coverage and is excluded from PSMC, MSMC, and SMC++ analysis. The other US sample has 34× coverage, but interestingly its *N*_e_ plot is similar to that of *T. aduncus* instead of East Asian *T. truncatus.* This observation is likely genuine rather than artifactual, because even a variation in coverage from 10× to 43× among East Asian samples does not affect the result.

To evaluate the robustness of the results to different parameters of mutation rate and generation time, we scaled the PSMC results for both species at generation times of 14, 21, and 28 years and mutation rates of 1.5×10^−8^, 2.5×10^−8^, and 5×10^−8^ per site per year. The range of generation time was chosen based on realistic values across cetacean species (Taylor et al. 2007). Furthermore, we used 100 randomizations for each of the individuals in PSMC to obtain the confidence intervals. The contrasting patterns in *N*_e_ trends between the two species are valid for a wide range of parameters. The parameters required to generate similar *N*_e_ trends in the two species would be highly unrealistic.

To examine the potential role of population structure in generating the observed temporal trends in Ne, we performed the pseudo-diploid analysis (Li and Durbin 2011; Prado-Martinez et al. 2013). Haploid sequences were generated from each individual using the seqtk program provided by Heng Li (<u>https://github.com/lh3/seqtk</u>). Sites with low quality were excluded using the flag −q 20, and one of the alleles was randomly chosen at heterozygous sites using the flag - *r*. The haploid genomes from two individuals obtained in the previous step were merged to create pseudo-diploid genomes, which was then subjected to the PSMC analysis.

While the PSMC analysis is able to infer the temporal trends of *N*_e_ based on the whole genome sequence of a single individual, the MSMC (Multiple Sequentially Markovian Coalescent) method is able to integrate information from multiple individuals of the same species to make more robust inferences (Schiffels and Durbin 2014). We used the script bamCaller.py provided along with the MSMC program to separately identify SNPs from each dolphin species. Statistical phasing of the SNP calls by the program shapeit requires at least 10 individuals from each population when a SNP reference panel is unavailable. However, we do not have a reference panel. While a previous study (Zhou et al. 2017) phased SNP data with only four individuals, they had access to a recombination rate map, which is not available for the dolphins. Hence, we converted our data to MSMC input format without phasing, using the script generate_multihetsep.py. The MSMC program was run with the option --fixedRecombination for each species separately. We did not study population divergences using the relative gene flow analysis because using unphased data can bias this analysis (Schiffels and Durbin 2014).

The MSMC method is better suited for data in which the phases of the genotype calls are known. However, in our case, accurate phasing is not possible due to the lack of a large number of samples. In such cases, the SMC++ method is an excellent alternative to the MSMC analysis. In addition to the genomic distribution of variable sites, SMC++ also utilizes information from the site frequency spectrum across individuals and has been shown to perform more reliably than MSMC, especially when using unphased data (Terhorst et al. 2016). In addition to not requiring phased data, SMC++ is also able to infer split times in diverged populations. The VCF (Variant Call Format) file with genotype information for all sites, including non-variant sites was generated using samtools. This file was converted to SMC format using the vcf2smc option in SMC++ program for all individuals of each population separately. The repeat annotation and numt regions identified in the genome were passed as the mask file to *vcf2smc* command with the −m flag. Population size histories were estimated using the *estimate* option in SMC++ with a mutation rate of 1.21×10^−9^ per site per year for both dolphin species (Taylor et al. 2007; Yim et al. 2014). Because the ancestral state could not be reliably reconstructed for our focal species, we used only the folded frequency spectrum option while running SMC++. Documentation for SMC++ suggests various parameters that need to be experimented with to identify the settings best suited for each dataset. With the default settings, our dataset was not generating results for more recent times. Hence, we set the option --t1 to 10 generations to extend the results to recent times. However, this resulted in over-fitting of the curves as well as too much oscillation. To correct this issue, we decreased the --regularization-penalty to a value of 5 based on recommendations in the SMC++ documentation. The parameter for thinning was set at 3000 and —ftol was increased to 0.01. With these settings, we were able to extend the estimates of population size by SMC++ to recent times.

We next inferred the split times for the species pair using the split option of SMC++ after estimating the joint frequency spectrum for both species with the same parameter settings as described in the previous paragraph. The split option of SMC++ provides an independent validation of the results from the pseudo-diploid analysis from PSMC. The latest version of SMC++ (version 1.11.1) does not work with data from single individuals because the site frequency estimates from single individuals are unreliable. Hence, we grouped two randomly selected individuals of *T. truncatus* (from the East Asian population) into population 1 and two other individuals of *T. truncatus* (from the East Asian population) into population 2. These two populations were used to infer the split times between *T. truncatus* individuals. Similarly, *T. aduncus* individuals were randomly grouped into two populations and used to estimate the split times among *T. aduncus* individuals.

The population structure of all sampled individuals was investigated using Principal Component Analysis (PCA) with the software ngsTools (Fumagalli et al. 2014). First, the posterior probability of genotypes was calculated using ANGSD (Korneliussen et al. 2014). These posterior probability values were then used to estimate the covariance matrix between individuals using the program ngsCovar. Finally, this covariance matrix was decomposed into eigenvectors using R to obtain the principal components. Similarly, we used the -doIBS option in ANGSD to sample a single base and calculate pairwise distances between each pair of individuals. This pair-wise distance matrix was scaled to obtain a multidimensional scaling plot.

### Population genetic statistics

The single nucleotide variants at bi-allelic sites that had no missing data in any of the individuals were used to estimate population genetic statistics such as Fst (differentiation), dxy (divergence), and π (nucleotide diversity) using Heirfstat (Goudet 2005). We further used ANGSD (Korneliussen et al. 2014) and ngsTools (Fumagalli et al. 2014) to estimate population genetic summary statistics. The values estimated from the two methods were in agreement.

## ACKNOWLEDGMENTS

We thank researchers who provided public access to sequencing data through the Short Read Archive, Yun Song and Jonathon Terhorst for advice on SMC++ analysis, and Andy Foote for valuable comments on an earlier draft. Byung-Yeob Kim provided some *T. aduncus* tissue samples. This research was supported by a grant from the Collaborative Genome Program (20140428) funded by the Ministry of Oceans and Fisheries, Korea and a National Research Foundation of Korea (NRF) grant from the Korean government (MSIP) (NRF-2015R1A4A1041997) to JKP. J.Z. was supported by U.S. National Institutes of Health research grant R01GM103232.

## Legends for supplementary figures

**Fig. S1**. Robustness of the inferred contrasting demographic changes of two dolphin species in the last glacial. (A) PSMC estimates of effective population size (Ne) over time inferred from the autosome scaffolds of one representative *T. truncatus* individual and one representative *T. aduncus* individual. Thick lines represent the median and thin light lines of the same color correspond to 100 bootstrap samples. (B) The PSMC analysis were conducted at different combinations of generation time (g = 14, 21, or 28 years) and mutation rate (μ = 1.5×10^−8^, 2.5×10^−8^, or 5.0×10^−8^ per site per generation). (C) PSMC analysis without making assumptions about the mutation rate and generation time.

**Fig. S2**. Analysis of population structure across the sampled dolphins by (A) multidimensional scaling and (B) principal component analysis. Red dots indicate *T. aduncus,* while blue dots indicate *T. truncatus.* Two *T. truncatus* individuals from Penglai, China are very close on these diagrams and carry the same label.

**Fig. S3**. PSMC estimates of *N*_e_ over time in the *T. truncatus* individual from Mississippi Sound, Louisiana.

**Fig. S4**. Population split time analysis of *T. truncatus* and *T. aduncus* using joint frequency spectrums. Red lines indicate *T. aduncus*, while blue lines indicate *T. truncatus*. The solid and dashed lines respectively represent trajectories estimated using two randomly selected individuals and the other two individuals from the four (East Asian) individuals sampled per species. The points at which the trajectories of the same color diverge are the split times estimated by SMC++.

**Fig. S5**. Pseudo-diploid PSMC analysis of *T. truncatus.* Dashed lines show pseudo-diploids between East Asian individuals (yellow) and those between Louisiana and East Asian individuals (red).

